# Mitochondrial One-Carbon Metabolism is Required for TGF-β-Induced Glycine Synthesis and Collagen Protein Production

**DOI:** 10.1101/2023.11.07.566074

**Authors:** Angelo Y. Meliton, Rengül Cetin-Atalay, Yufeng Tian, Jennifer C. Houpy Szafran, Kun Woo D. Shin, Takugo Cho, Kaitlyn A. Sun, Parker S. Woods, Obada R. Shamaa, Bohao Chen, Alexander Muir, Gökhan M. Mutlu, Robert B. Hamanaka

## Abstract

A hallmark of Idiopathic Pulmonary Fibrosis is the TGF-β-dependent activation of lung fibroblasts, leading to excessive deposition of collagen proteins and progressive scarring. We have previously shown that synthesis of collagen by lung fibroblasts requires *de novo* synthesis of glycine, the most abundant amino acid in collagen protein. TGF-β upregulates the expression of the enzymes of the *de novo* serine/glycine synthesis pathway in lung fibroblasts through mTORC1 and ATF4- dependent transcriptional programs. SHMT2, the final enzyme of the *de novo* serine/glycine synthesis pathway, transfers a one-carbon unit from serine to tetrahydrofolate (THF), producing glycine and 5,10-methylene-THF (meTHF). meTHF is converted back to THF in the mitochondrial one-carbon (1C) pathway through the sequential actions of MTHFD2 (which converts meTHF to 10-formyl-THF), and either MTHFD1L, which produces formate, or ALDH1L2, which produces CO_2_. It is unknown how the mitochondrial 1C pathway contributes to glycine biosynthesis or collagen protein production in fibroblasts, or fibrosis *in vivo*. Here, we demonstrate that TGF-β induces the expression of *MTHFD2*, *MTHFD1L*, and *ALDH1L2* in human lung fibroblasts. *MTHFD2* expression was required for TGF-β-induced cellular glycine accumulation and collagen protein production. Combined knockdown of both *MTHFD1L* and *ALDH1L2* also inhibited glycine accumulation and collagen protein production downstream of TGF-β; however knockdown of either protein alone had no inhibitory effect, suggesting that lung fibroblasts can utilize either enzyme to regenerate THF. Pharmacologic inhibition of MTHFD2 recapitulated the effects of *MTHFD2* knockdown in lung fibroblasts and ameliorated fibrotic responses after intratracheal bleomycin instillation *in vivo*. Our results provide insight into the metabolic requirements of lung fibroblasts and provide support for continued development of MTHFD2 inhibitors for the treatment of IPF and other fibrotic diseases.

## INTRODUCTION

Idiopathic Pulmonary Fibrosis (IPF) is a progressive, fatal disease, which has a median survival of 3.5 years and affects approximately 150,000 people in the United States (1, 2). IPF is characterized by the Transforming Growth Factor-β (TGF-β)-dependent activation of lung fibroblasts, leading to the excessive secretion of extracellular matrix proteins, including collagens (3–6). Replacement of healthy lung tissue with matrix proteins leads to progressive loss of lung function and thus, the production of collagens by lung fibroblasts represents an important therapeutic target for the treatment of IPF (6–8).

Metabolic reprogramming during the process of fibroblast activation is an emerging hallmark of IPF and an increasingly-studied target for therapeutic intervention (9, 10). We and others have demonstrated that *de novo* synthesis of glycine, the most abundant amino acid in collagen protein is required for production of collagen downstream of TGF-β (11–14). This requires the mTOR and ATF4-dependent induction of the enzymes of the *de novo* serine/glycine synthesis pathway, which converts the glycolytic intermediate 3-phosphoglycerate into the amino acids serine and glycine (13, 15). Inhibition of this pathway reduces TGF-β-induced collagen protein production *in vitro*, and ameliorates bleomycin-induced lung fibrosis *in vivo* (11, 12, 16).

The last enzyme of the *de novo* serine/glycine synthesis pathway, serine hydroxymethyltransferase 2 (SHMT2), is an entry point for carbon into the mitochondrial one-carbon (1C) pathway (17–22). SHMT2 catalyzes the transfer of a 1C group from serine to tetrahydrofolate (THF), producing glycine and 5,10-methylene-THF (meTHF). In order for the SHMT2 reaction to proceed, mitochondrial meTHF must be recycled back to THF by the enzymes of the mitochondrial 1C pathway. First, the bifunctional dehydrogenase/cyclohydrolase, MTHFD2 (meTHF Dehydrogenase 2) catalyzes the conversion of meTHF to 10-formyl-THF. 10-formyl-THF is then converted back to THF by either MTHFD1L (meTHF Dehydrogenase 1 Like), producing formate, or by ALDH1L2 (Aldehyde Dehydrogenase Family 1 Member L2), producing CO_2_ and reducing NADP^+^ to NADPH. It is unknown how 1C metabolism contributes to glycine synthesis and collagen protein production downstream of TGF-β signaling.

Here, we demonstrate that TGF-β induces the expression of *MTHFD2*, *MTHFD1L*, and *ALDH1L2* in an mTORC1 and ATF4-dependent manner. MTHFD2 is required for intracellular glycine accumulation downstream of TGF-β and for collagen protein production in lung fibroblasts. The combined knockdown of *MTHFD1L* and *ALDH1L2* recapitulated the effects of *MTHFD2* knockdown; however, the individual knockdown of either enzyme did not affect glycine or collagen accumulation, suggesting that lung fibroblasts have flexibility to utilize either enzyme to regenerate THF. We further show that the inhibition of MTHFD2 using DS18561882 recapitulates the effects of *MTHFD2* knockdown on glycine and collagen production downstream of TGF-β. Furthermore, treatment of mice with DS18561882 ameliorates fibrotic responses after intratracheal instillation of bleomycin. Our results demonstrate that MTHFD2 may be a viable therapeutic target for IPF and other fibrotic diseases.

## RESULTS

### TGF-β induces the expression of mitochondrial 1C pathway enzymes through mTORC1 and ATF4

We have previously demonstrated that lung fibroblasts depend on *de novo* synthesis of glycine to support production of collagen protein downstream of TGF-β (11, 12). Synthesis of glycine from serine produces 1C units which are transferred to THF, producing 5,10-methylene-THF (meTHF), which must be recycled back to THF to support glycine production by SHMT2 **(Fig. 1A)** (22). To determine how the TGF-β regulates the expression of 1C enzymes, we treated normal human lung fibroblasts (NHLFs) with TGF-β (1ng/mL) and measured mRNA and protein expression of the mitochondrial 1C enzymes MTHFD2, ALDH1L2, and MTHFD1L **(Fig. 1B)**. TGF-β significantly increased the mRNA and protein expression of MTHFD2 and ALDH1L2, with a smaller increase in MTHFD1L expression **(Fig. 1B, 1C, S1A-C)**.

**Figure 1.**
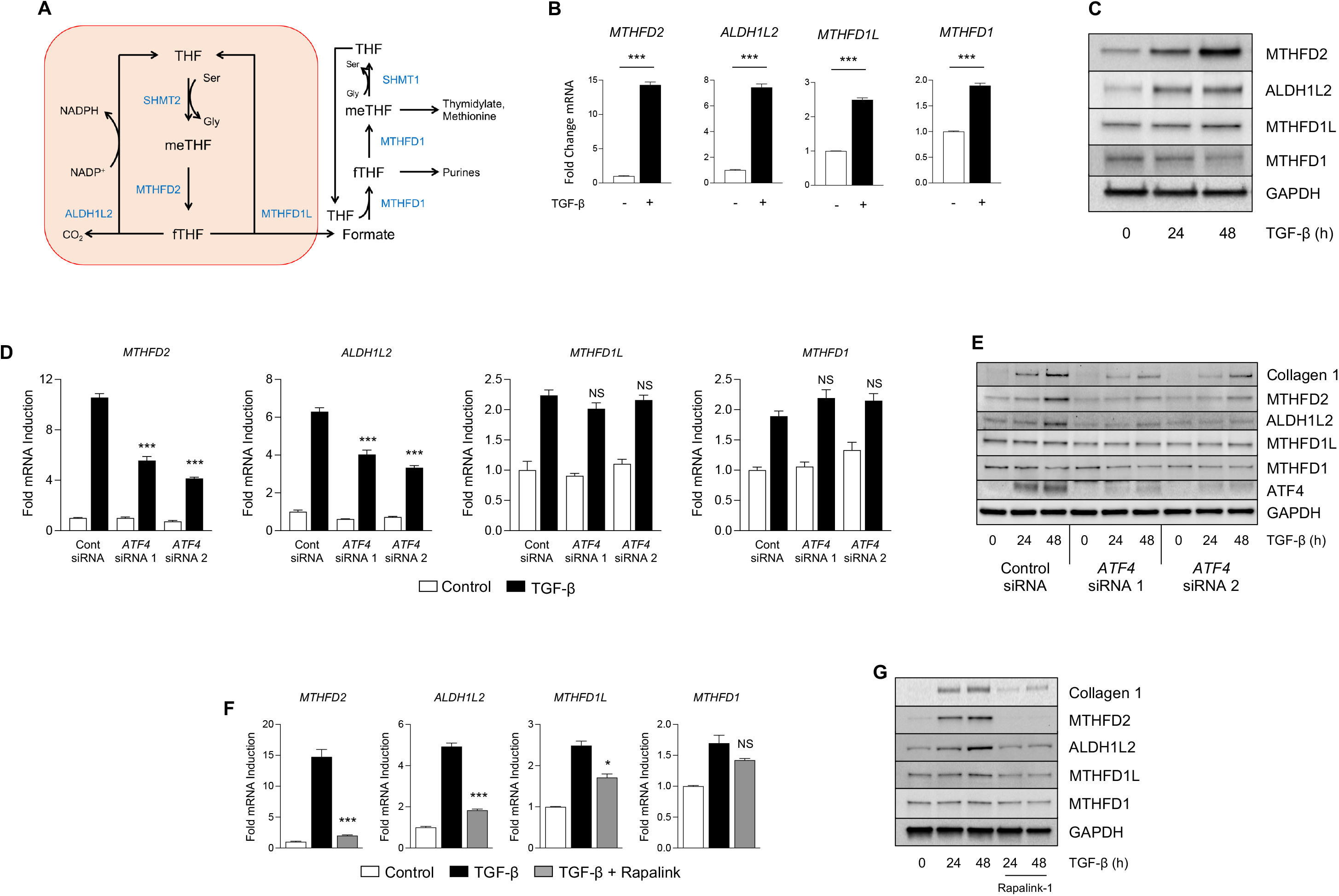
Mitochondrial one-carbon enzymes are induced by TGF-β. **(A)** Schematic representation of metabolite flux through the mitochondrial and cytosolic one-carbon pathways. **(B)** qRT-PCR analysis of 1C enzyme mRNA expression in normal human lung fibroblasts (NHLFs) cultured in the presence or absence of TGF-β for 24 hours. **(C)** Western blot analysis of 1C enzyme protein expression in NHLFs treated with TGF-β for the indicated intervals. **(D)** qRT-PCR analysis of 1C enzyme mRNA expression in NHLFs transfected with siRNA targeting *ATF4* or nontargeting siRNA. Cells were treated with TGF-β or left untreated for 24 hours. **(E)** Western blot analysis of 1C enzyme protein expression in NHLFs transfected with siRNA targeting *ATF4* or nontargeting siRNA. Cells were treated with TGF-β for the indicated intervals. **(F)** qRT-PCR analysis of 1C enzyme mRNA expression in NHLFs either left untreated or treated with TGF-β in the presence or absence of the mTORC1 inhibitor Rapalink-1. **(G)** Western blot analysis of 1C enzyme protein expression in NHLFs treated with TGF-β for the indicated intervals in the presence or absence of Rapalink-1. Bar graphs represent mean ± SEM, n=3. \**P*<0.05, \*\**P*<0.01, \*\*\**P*<0.001.

Flux through the mitochondrial 1C pathway has been shown to be required to supply 1C units to the cytosolic 1C pathway through formate production and mitochondrial export. Formate overflow from the mitochondria supplies 1C units for generation of purines, thymidylate, and methionine **(Fig. 1A)** (22). In contrast to the mitochondrial pathway, which uses multiple enzymes, the cytoplasmic 1C pathway is catalyzed only by MTHFD1. We found that while *MTHFD1* transcript levels were increased by TGF-β, its protein expression was reduced compared with untreated cells **(Fig. 1B, 1C, S1D)**.

We have previously demonstrated that induction of the serine/glycine synthesis pathway downstream of TGF-β is dependent on mTORC1 and ATF4 (15). Thus, we sought to determine the requirement of these factors for the regulation of 1C enzyme expression downstream of TGF- β. We found that TGF-β-mediated induction of *MTHFD2* and *ALDH1L2* was inhibited by *ATF4* knockdown, while *MTHFD1L* and *MTHFD1* were unaffected **(Figure 1D, 1E, S1E-H)**. Consistent with a role of mTORC1 promoting ATF4 activation, inhibition of mTORC1 with the selective inhibitor Rapalink-1 resulted in similar reduced expression of *MTHFD2* and *ALDH1L2*, with a partial inhibition of *MTHFD1L* induction downstream of TGF-β **(Fig. 1F, 1G, S1I-L)**.

### Mitochondrial 1C metabolism is required for TGF-β-induced collagen protein production in NHLFs

We have previously shown that SHMT2-mediated production of glycine is required to support collagen protein production in lung fibroblasts (11). Because the SHMT reaction requires mitochondrial THF regeneration **(Fig. 1A)**, we sought to determine whether mitochondrial 1C enzymes are required for collagen protein production downstream of TGF-β. Consistent with the specific importance of mitochondrial 1C metabolism for collagen production, knockdown of *MTHFD2* was sufficient to inhibit TGF-β-induced collagen protein production **(Fig 2A, 2B)**, while knockdown of the cytosolic enzyme, MTHFD1 did not affect collagen induction downstream of TGF-β **(Fig. 2C, 2D)**. To determine whether the enzymes downstream of MTHFD2 are required TGF-β-induced collagen production, we knocked down either *ALDH1L2* **(Fig. 2E, 2F)**, *MTHFD1L* **(Fig. 2G, 2H)**, or both *ALDH1L2* and *MTHFD1L* **(Fig. 2I, 2J)**. Surprisingly, knockdown of either *ALDH1L2* or *MTHFD1L* did not affect collagen induction in NHLFs treated with TGF-β; however, the combined knockdown of both enzymes did reduce collagen induction, suggesting that while MTHFD2 is required for collagen production downstream of TGF-β, NHLFs have flexibility to use either ALDH1L2 or MTHFD1L in order to regenerate mitochondrial THF.

**Figure 2.**
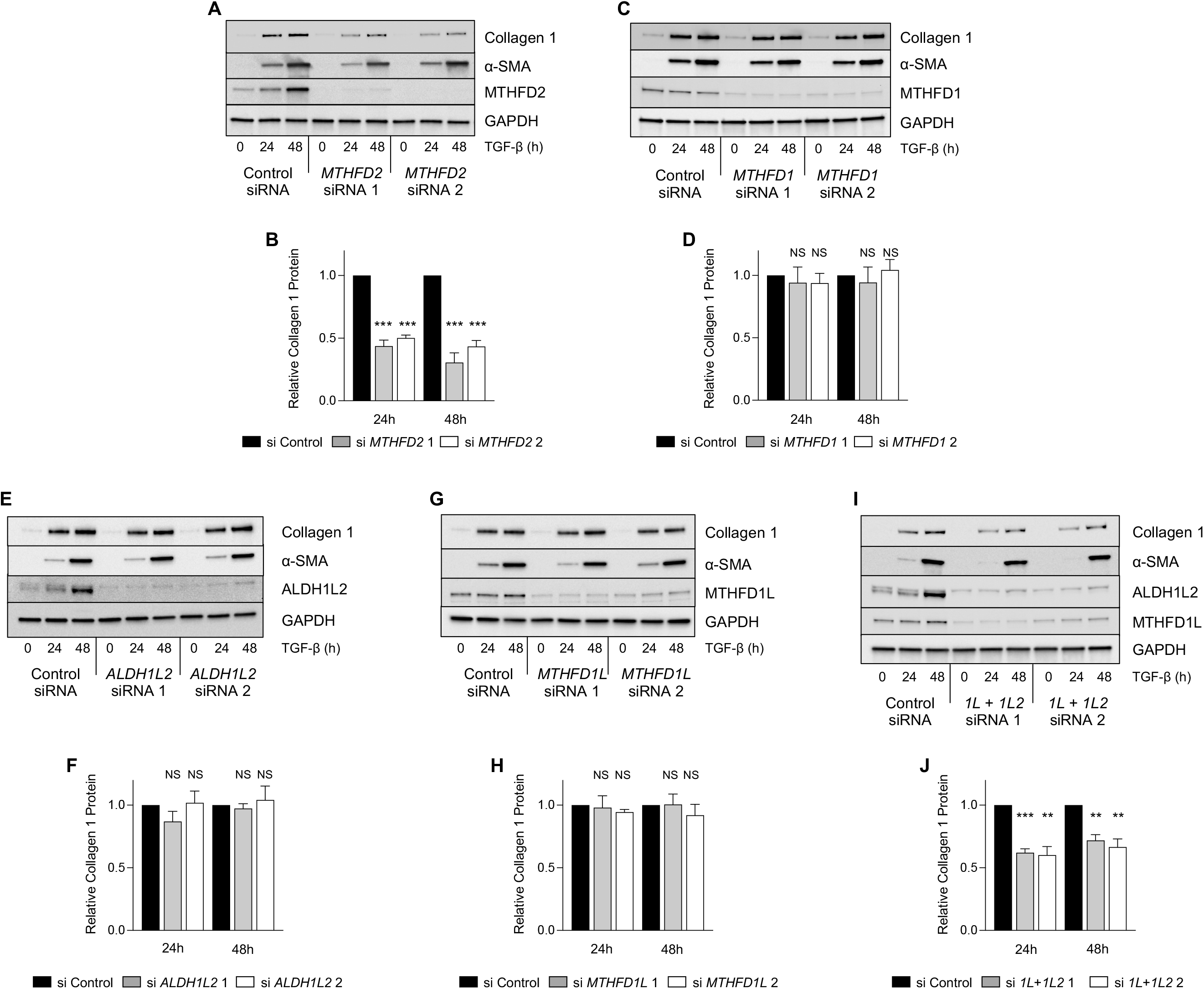
Mitochondrial one-carbon metabolism is required for TGF-β-induced collagen protein production. **(A)** Western blot analysis of collagen and α-smooth muscle actin (α-SMA) protein expression in normal human lung fibroblasts (NHLFs) transfected with siRNA targeting *MTHFD2* or nontargeting siRNA. Cells were treated with TGF-β for the indicated intervals. **(B)** Quantification of relative Collagen 1 levels in *MTHFD2* knockdown NHLFs at the indicated intervals. **(C)** Western blot analysis of collagen and α-SMA protein expression in NHLFs transfected with siRNA targeting *MTHFD1* or nontargeting siRNA. Cells were treated with TGF-β for the indicated intervals. **(D)** Quantification of relative Collagen 1 levels in *MTHFD1* knockdown NHLFs at the indicated intervals. **(E)** Western blot analysis of collagen and α-SMA protein expression in NHLFs transfected with siRNA targeting *ALDH1L2* or nontargeting siRNA. Cells were treated with TGF-β for the indicated intervals. **(F)** Quantification of relative Collagen 1 levels in *ALDH1L2* knockdown NHLFs at the indicated intervals. **(G)** Western blot analysis of collagen and α-SMA protein expression in NHLFs transfected with siRNA targeting *MTHFD1L* or nontargeting siRNA. Cells were treated with TGF-β for the indicated intervals. **(H)** Quantification of relative Collagen 1 levels in *MTHFD1L* knockdown NHLFs at the indicated intervals. **(I)** Western blot analysis of collagen and α-SMA protein expression in NHLFs transfected with siRNA targeting both *ALDH1L2* and *MTHFD1L* or nontargeting siRNA. Cells were treated with TGF-β for the indicated intervals. **(J)** Quantification of relative Collagen 1 levels in *ALDH1L2*/*MTHFD1L* knockdown NHLFs at the indicated intervals. Bar graphs represent mean ± SEM, n=3. \**P*<0.05, \*\**P*<0.01, \*\*\**P*<0.001.

Our results suggest that mitochondrial formate production and cytosolic 1C metabolism are dispensable for collagen protein production downstream of TGF-β. Consistent with this, formate supplementation was insufficient to rescue the effects of *MTHFD2* knockdown on collagen production by NHLFs **(Fig. S2A, S2B)**. Furthermore, treatment of NHLFs with methotrexate, which reduces cytosolic THF by inhibiting dihydrofolate reductase, had no effect on collagen protein production downstream of TGF-β **(Fig. S2C)**. Finally, to determine whether folate supplementation into the media could rescue *MTHFD2* knockdown, we added increasing doses of 5-methyl-THF to the media of *MTHFD2* knockdown NHLFs. Consistent with an absolute requirement of mitochondrial THF regeneration for collagen protein production by lung fibroblasts, 5-methyl-THF was unable to rescue the effects of *MTHFD2* knockdown **(Fig. S2D-S2E)**. This finding is consistent with previous reports demonstrating high levels of compartmentalization of cellular folate pools with limited transport (23, 24).

### Mitochondrial 1C metabolism is required for TGF-β-induced cellular glycine accumulation

To determine how mitochondrial 1C metabolism contributes to *de novo* glycine synthesis in NHLFs, we measured cellular serine and glycine levels in control and *MTHFD2* knockdown cells using gas chromatography mass spectrometry (GC-MS). Cells were cultured in media containing ^13^C2-glycine (glycine labeled with ^13^C on both carbon atoms) to allow us to determine the relative contribution of extracellular (M+2) glycine as well as glycine synthesized *de novo* from either extracellular glucose or serine (M+0) to total cellular glycine pools **(Fig. S3A)**. As shown in **Figure S3B**, unlabeled glycine, derived from both extracellular glucose and extracellular serine constituted over half of intracellular glycine in untreated cells. TGF-β increased both total cellular glycine and levels of *de novo*-synthesized (M+0) glycine in NHLFs **(Fig. S3C, S3D)**. Both total glycine and *de novo*-synthesized glycine were reduced in *MTHFD2* knockdown cells **(Fig. S3C, S3D)**. MTHFD2 knockdown was also associated with a significant increase in cellular serine levels as well as increased levels of cellular serine labeled from extracellular glycine **(Fig. S3E-G)**. This suggests that in the absence of MTHFD2, catabolism of serine is inhibited and reverse flux through SHMT2 is increased, leading to increased serine synthesis from glycine. This was confirmed in *SHMT2* knockdown cells which also displayed reduced total glycine and *de novo* synthesized glycine levels **(Fig. S4A-C)**. While *SHMT2* knockdown cells had elevated total serine levels compared with knockdown cells, labeling of serine from extracellular glycine was inhibited in *SHMT2* knockdown cells **(Fig. S4D-F)**.

To better parse the relative contributions of the individual mitochondrial 1C enzymes to cellular glycine production, we labeled control and *MTHFD2* knockdown cells with 2,3,3-D3-Serine (serine labeled with one deuterium atom on the 2-carbon, and 2 deuterium atoms on the 3-carbon), which has been used to demonstrate the contribution of 1C metabolism to cellular NADPH homeostasis (25, 26) **(Fig. 3A)**. Consistent with our findings in **Figure S3**, cellular glycine levels, and glycine synthesized from serine were increased in TGF-β-treated cells **(Fig. 3B, 3C, S5A)**. Total cellular serine levels were elevated in *MTHFD2* knockdown cells, as were M+1 and M+2 serine resulting from reverse flux through SHMT **(Fig. 3D, 3E, S5B)**. The activity of the pathways downstream of MTHFD2 can be measured by tracing the presence of deuterium atoms into proline. As has been previously demonstrated, the reduction of NADP^+^ by MTHFD2 and ALDH1L2 leads to labeling of mitochondrial NADPH with deuterium, which can then be transferred to proline during its *de novo* synthesis (25, 26). Indeed, we found that TGF-β increased cellular proline levels and labeling of cellular proline (M+1) with deuterium from serine **(Fig. 3F, 3G, S5C)**. While labeling of proline was abolished in *MTHFD2* knockdowns, TGF-β-induced proline accumulation was not inhibited, demonstrating that while mitochondrial 1C metabolism contributes to mitochondrial NADPH pools, it is not required to support cellular proline production. Formate, which is exported from mitochondria and can be released from the cell, was elevated in the media of control cells after TGF-β treatment, consistent with increased MTHFD1L activity **(Fig. 3H)**. *MTHFD2* knockdown reduced formate accumulation in the media both in control and TGF-β-treated NHLFs.

**Figure 3.**
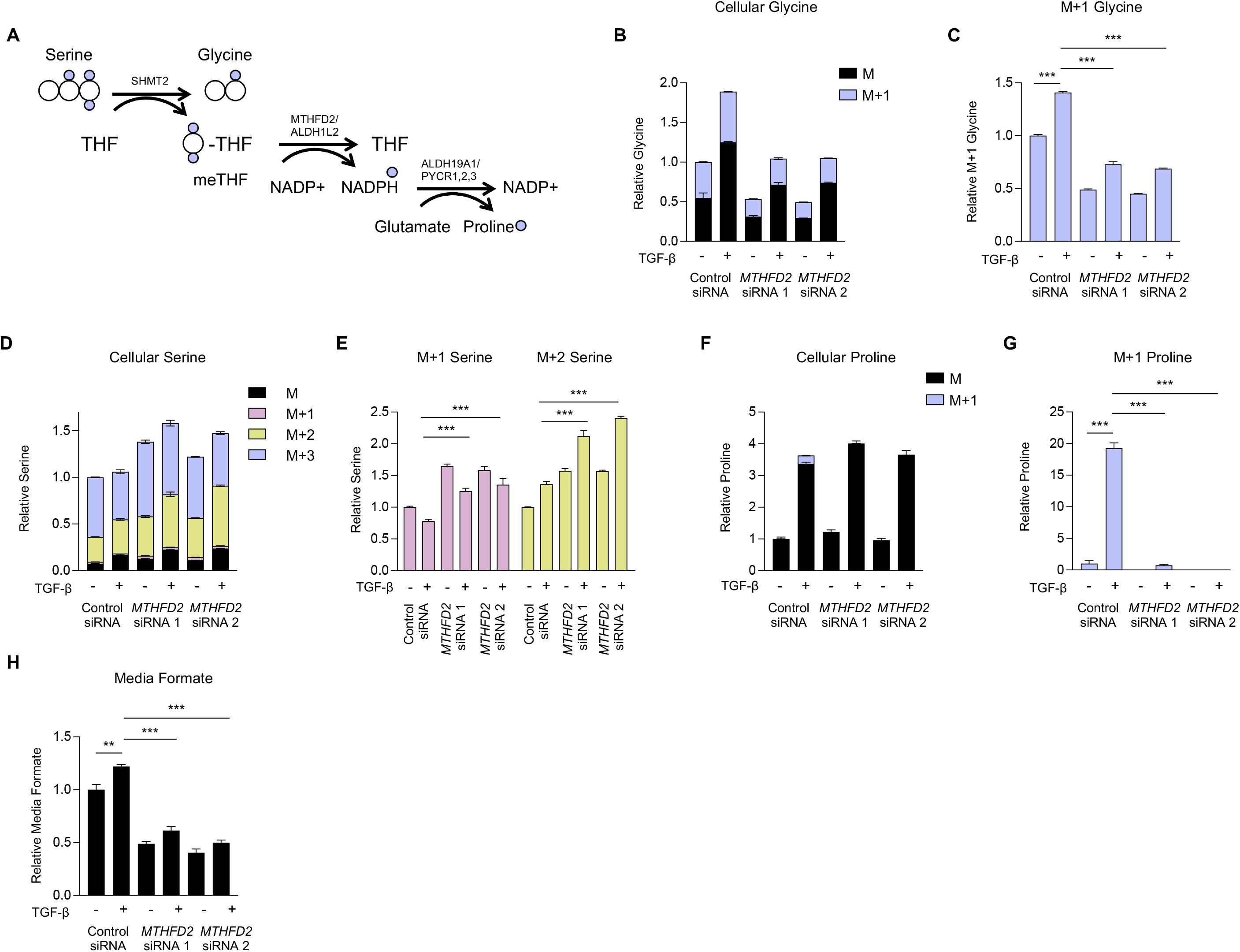
MTHFD2 is required for increased cellular glycine levels downstream of TGF-β. **(A)** Schematic representation of metabolite labeling downstream of 2,3,3-D3-Serine. Normal human lung fibroblasts (NHLFs) were transfected with siRNA targeting *MTHFD2* or nontargeting siRNA. Cells were labeled with 2,3,3-D3-Serine and treated with TGF-β for 48 hours or left untreated. **(B)** Analysis of cellular glycine after labeling with 2,3,3-D3-Serine in NHLFs treated with TGF-β or left untreated. **(C)** Relative levels of M+1 glycine from (B). **(D)** Analysis of cellular serine after labeling with 2,3,3-D3-Serine in NHLFs treated with TGF-β or left untreated. **(E)** Relative levels of M+1 and M+2 serine from (D). **(F)** Analysis of cellular proline after labeling with 2,3,3-D3-Serine in NHLFs treated with TGF-β or left untreated. **(G)** Relative levels of M+1 and proline from (F). **(H)** Relative levels of media formate content in NHLFs treated with TGF-β or left untreated. Bar graphs represent mean ± SEM, n=3. \**P*<0.05, \*\**P*<0.01, \*\*\**P*<0.001.

### Lung fibroblasts can use either ALDH1L2 or MTHFD1L to support glycine production

Consistent with a lack of effect of *ALDH1L2* or *MTHFD1L* knockdown on TGF-β-induced collagen production, knockdown of either protein did not prevent TGF-β-induced increases in cellular glycine levels **(Fig. 4A, 4E, S6A, S7A)** or lead to a significant decrease in glycine derived from 2,3,3-D3-Serine (M+1) **(Fig. S6B, S7B)**. Interestingly, an increase in total serine levels **(Fig 4B, 4F, S6C, S7C)** and in M+2 serine **(Fig. S6D, S7D)** was detected after either *ALDH1L2* or *MTHFD1L* knockdown, suggesting a level of inhibition in the mitochondrial one carbon pathway; however, this was insufficient to inhibit TGF-β-induced glycine accumulation. Consistent with the reduction of NADP^+^ by ALDH1L2, TGF-β-induced proline labeling was abolished in *ALDH1L2* knockdowns **(Fig. 4C, S6F)** while *MTHFD1L* knockdown had no effect **(Fig 4G, S7F)**. Total proline accumulation was unaffected by either knockdown, consistent with our findings from MTHFD2 knockdowns **(Fig. S6E, S7E)**. Consistent with a role for MTHFD1L in producing formate downstream of MTHFD2, *MTHFD1L* knockdown inhibited formate accumulation in the media after TGF-β treatment **(Fig. 4H)**, while *ALDH1L2* knockdown had no effect **(Fig 4D)**. Our results suggest that while MTHFD2 is essential for elevation of cellular glycine levels downstream of TGF-β, NHLFs have the flexibility to utilize either ALDH1L2 or MTHFD1L to regenerate mitochondrial THF. Consistent with this, combined knockdown of both *MTHFD1L* and *ALDH1L2*, inhibited TGF-β-induced production of glycine from serine **(Fig. 4I, S8B)** and mimicked the effect of *MTHFD2* knockdown on serine and proline labeling after TGF-β treatment as well as on media formate accumulation **(Fig. 4J-4L, S8)**. Consistent with a lack of effect of *MTHFD1* knockdown on collagen protein production, *MTHFD1* knockdown did not affect accumulation or labeling of glycine, serine, or proline downstream of TGF-β **(Fig. S9)**.

**Figure 4.**
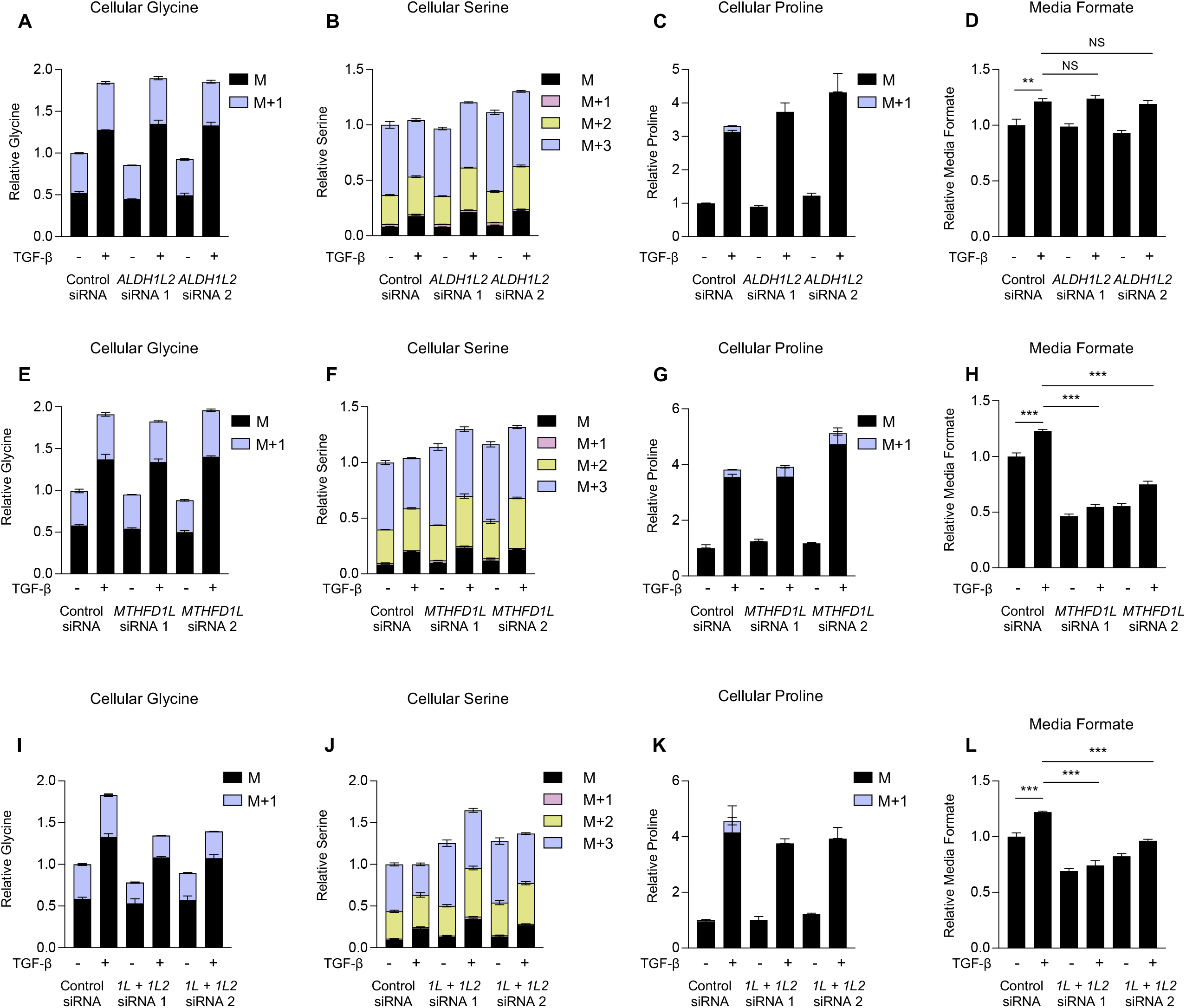
Human lung fibroblasts can utilize either ALDH1L2 or MTHFD1L to support glycine synthesis. Normal human lung fibroblasts (NHLFs) were transfected with the indicated siRNA. Cells were labeled with 2,3,3-D3-Serine and treated with TGF-β for 48 hours or left untreated. **(A)** Analysis of cellular glycine in *ALDH1L2* knockdown NHLFs after labeling with 2,3,3-D3-Serine in the presence or absence of TGF-β. **(B)** Analysis of cellular serine in *ALDH1L2* knockdown NHLFs after labeling with 2,3,3-D3-Serine in the presence or absence of TGF-β. **(C)** Analysis of cellular proline in *ALDH1L2* knockdown NHLFs after labeling with 2,3,3-D3-Serine in the presence or absence of TGF-β. **(D)** Relative levels of media formate content in *ALDH1L2* knockdown NHLFs treated with TGF-β or left untreated. **(E)** Analysis of cellular glycine in *MTHFD1L* knockdown NHLFs after labeling with 2,3,3-D3-Serine in the presence or absence of TGF-β. **(F)** Analysis of cellular serine in *MTHFD1L* knockdown NHLFs after labeling with 2,3,3- D3-Serine in the presence or absence of TGF-β. **(G)** Analysis of cellular proline in *MTHFD1L* knockdown NHLFs after labeling with 2,3,3-D3-Serine in the presence or absence of TGF-β. **(H)** Relative levels of media formate content in *MTHFD1L* knockdown NHLFs treated with TGF-β or left untreated. **(I)** Analysis of cellular glycine in *ALDH1L2*/*MTHFD1L* knockdown NHLFs after labeling with 2,3,3-D3-Serine in the presence or absence of TGF-β. **(J)** Analysis of cellular serine in *ALDH1L2*/*MTHFD1L* knockdown NHLFs after labeling with 2,3,3-D3-Serine in the presence or absence of TGF-β. **(K)** Analysis of cellular proline in *ALDH1L2*/*MTHFD1L* knockdown NHLFs after labeling with 2,3,3-D3-Serine in the presence or absence of TGF-β. **(L)** Relative levels of media formate content in *ALDH1L2*/*MTHFD1L* knockdown NHLFs treated with TGF-β or left untreated. Bar graphs represent mean ± SEM, n=3. \**P*<0.05, \*\**P*<0.01, \*\*\**P*<0.001.

### MTHFD2 regulates fibrotic responses *in vivo*

To determine whether mitochondrial one carbon metabolism might play a role in fibrotic processes *in vivo*, we examined human IPF lung tissue as well as lung tissue from mouse lung after induction of fibrosis with bleomycin. Immunohistochemical analysis demonstrated that MTHFD2 is highly expressed in fibrotic foci of IPF patients when compared with non-fibrotic areas of the same lung **(Fig. 5A)**. MTHFD2 was also highly induced in mouse lung tissue after bleomycin instillation **(Fig 5B)**. Using publicly available data sets of gene expression from IPF patients (27), we found that *MTHFD2* expression was negatively correlated with forced vital capacity (FVC), suggesting an involvement of MTHFD2 in disease progression **(Fig 5C)**.

**Figure 5.**
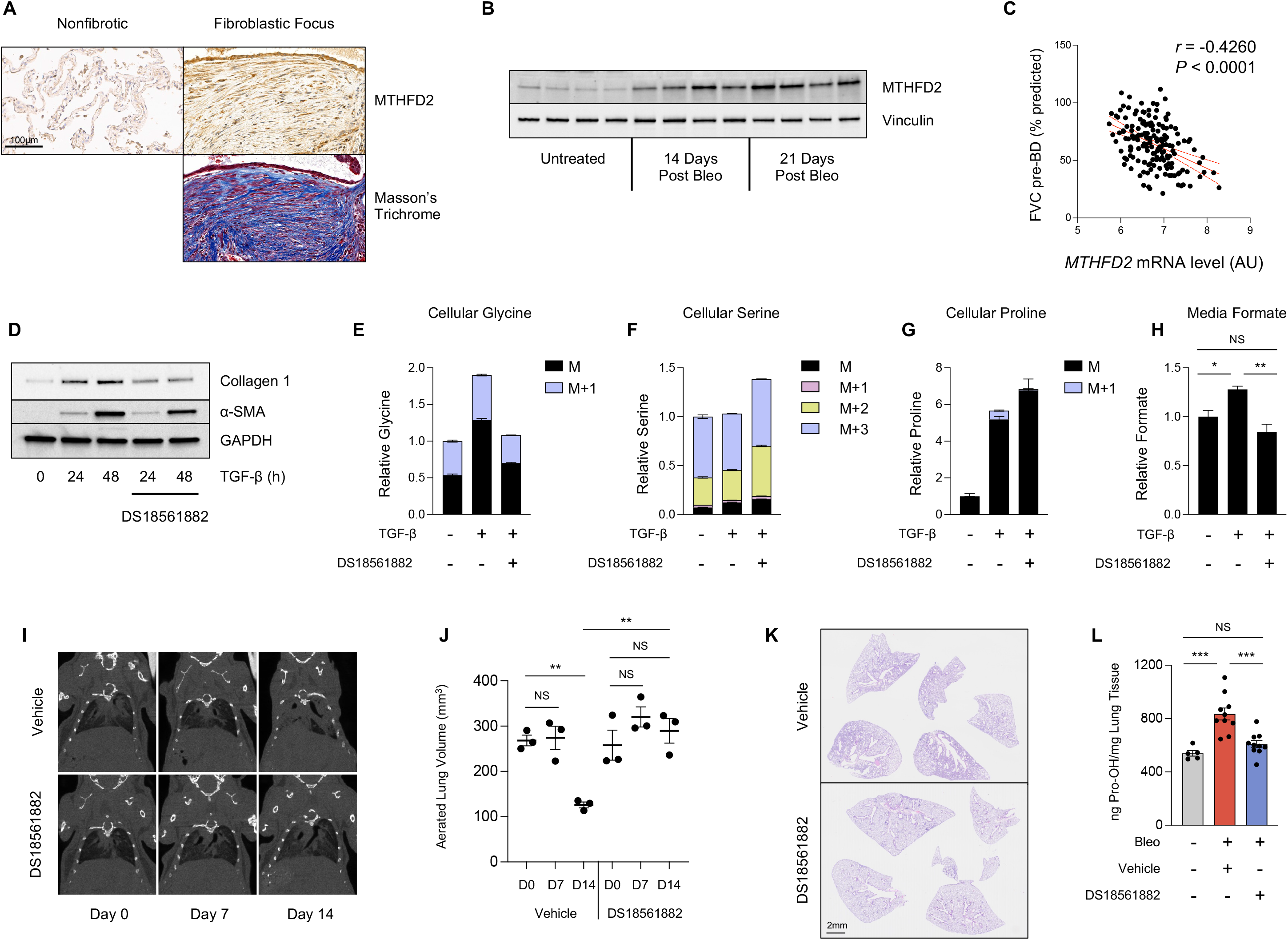
Pharmacologic inhibition of MTHFD2 inhibits fibrotic responses *in vivo*. **(A)** Histological analysis of MTHFD2 protein expression in fibrotic and nonfibrotic areas of paraffin-embedded lung sections from a patient with idiopathic pulmonary fibrosis. **(B)** Western blot analysis of MTHFD2 protein expression in mouse lung tissue either prior to, or 14 or 21 days after bleomycin instillation. **(C)** Pearson’s correlation of *MTHFD2* mRNA level and forced vital capacity before bronchodilator (FVC-pre-BD) as percentage of what was predicted for each patient. Data is from GSE32537. **(D)** Western blot analysis of collagen and α-smooth muscle actin protein expression in normal human lung fibroblasts (NHLFs) treated with TGF-β for the indicated intervals in the presence or absence of DS18561882 (5μM). **(E-G)** Analysis of cellular (E) glycine, (F) serine, and (G) proline in NHLFs after labeling with 2,3,3-D3-Serine in the presence or absence of TGF-β and DS18561882. **(H)** Relative levels of media formate content in NHLFs treated with TGF-β or left untreated in the presence or absence of DS18561882. **(I)** Representative coronal microCT images of mice prior to bleomycin instillation (Day 0), 7 days after bleomycin instillation (Day 7), and 14 days after bleomycin instillation (Day 14). Mice began receiving DS18561882 (125 mg/kg) or vehicle 8 days after bleomycin instillation. **(J)** Quantification of aerated lung volumes from microCT-scanned mice treated as in (I) (mean ± SEM). **(K)** Histological analysis of mouse lung tissue on day 21 after bleomycin instillation. Beginning on day 8, mice received daily IP injections of either DS18561882 or vehicle. **(L)** Hydroxyproline content of mouse lungs on day 21 after bleomycin or vehicle (saline) instillation. Beginning on day 8, mice received either DS18561882 or vehicle IP, daily (mean ± SEM). \**P*<0.05, \*\**P*<0.01, \*\*\**P*<0.001.

DS18561882 is a small molecule inhibitor of MTHFD2 that has been shown to inhibit cancer cell growth and T-cell-mediated inflammation *in vivo* (28–30). We found that treatment of NHLFs with DS18561882 inhibited TGF-β-induced collagen protein production **(Fig. 5D)**. Furthermore, DS18561882 treatment mimicked the effects of *MTHFD2* knockdown on cellular glycine accumulation after TGF-β and labeling from 2,3,3-D3-Serine **(Fig. 5E, S10A, S10B)**. Elevated cellular levels of serine, loss of labeling on proline from 2,3,3-D3-Serine, and inhibition of media formate accumulation were also observed in the presence of DS18561882, suggesting that the effects of DS18561882 are specific to MTHFD2 inhibition **(Fig. 5F-5H, S10)**.

We thus treated mice with DS18561882 to determine if MTHFD2 inhibition can therapeutically inhibit progression of lung fibrosis after bleomycin instillation. Mice were intratracheally instilled with bleomycin, which induces an acute lung injury/inflammatory state lasting approximately one week, followed by a fibrotic phase which lasts 2-3 weeks. To determine the therapeutic efficacy of MTHFD2 inhibition on active fibrotic processes, mice were treated with DS18561882 or vehicle beginning on day 8 after bleomycin instillation. Using micro computed tomography (microCT), we measured lung aeration at day 7 and day 14 after bleomycin instillation. All mice had similar levels of lung aeration prior to and at day 7 after bleomycin **(Fig 5I, 5J, S10G)**. Mice receiving vehicle displayed decreased aeration on day 14 concomitant with the induction of fibrosis. Mice receiving DS18561882 beginning on day 8 did not exhibit a significant drop in lung aeration on day 14. Endpoint analysis at day 21 post bleomycin revealed greatly reduced degree of fibrosis in DS18561882-treated mice **(Fig 5K, S10H)**. Lung hydroxyproline levels, an indicator of collagen content, were also significantly reduced in mice treated with DS18561882 **(Fig 5L)**. Together, our results suggest that pharmacologic targeting of MTHFD2 may be a viable therapeutic avenue for IPF.

## DISCUSSION

Metabolic reprogramming in lung fibroblasts is an emerging mechanism required to support matrix production and the development of fibrotic disease. We have previously demonstrated that TGF- β induces metabolic reprogramming in lung fibroblasts characterized by increased levels of *de novo* serine and glycine synthesis (11, 12, 15). Glycine constitutes one third of the primary structure of collagens, and we and others have found that *de novo* glycine synthesis supports collagen synthesis downstream of TGF-β (11–16). Because glycine synthesis by SHMT2 is linked with the mitochondrial one-carbon pathway, we sought to determine the roles and requirements of 1C metabolism in glycine synthesis and collagen protein production downstream of TGF-β.

We found that TGF-β significantly increases the expression of mitochondrial 1C enzymes in lung fibroblasts. Consistent with our previous findings that the enzymes of the serine/glycine synthesis pathway are regulated by mTORC1 and ATF4, we found that *MTHFD2* and *ALDH1L2* are induced by TGF-β through these same mechanisms. This is consistent with previous reports of regulation of these enzymes by mTORC1 and ATF4 downstream of insulin signaling (31–33). *MTHFD1L* has been shown to be regulated by transcription factors including NRF2 and c-myc (34–36). While the mechanism of *MTHFD1L* regulation by TGF-β remains to be determined, it is clear that metabolic pathways that support amino acid biosynthesis are among the primary targets of TGF-β-mediated transcriptional responses.

Our results demonstrate that MTHFD2 is a critical regulator of glycine synthesis and collagen protein production in lung fibroblasts. While these cells appear to have the flexibility to use either MTHFD1L or ALDH1L2 to regenerate mitochondrial THF for the SHMT2 reaction, MTHFD2 is essential for mitochondrial THF regeneration, and inhibition of MTHFD2 results in reduced cellular glycine levels and collagen protein production. Our results suggest that loss of MTHFD2 function not only inhibits *de novo* glycine synthesis, but promotes the reversal of the SHMT2 reaction, potentially making MTHFD2 a more effective therapeutic target for fibrotic disease. While we cannot rule out potential additional roles for formate production by MTHFD1L and activity of the cytoplasmic 1C pathway in lung fibroblasts, our findings demonstrate that these are dispensable for collagen protein production. The cytoplasmic 1C pathway has been shown to reverse flux after inhibition of the mitochondrial one-carbon pathway (25), and may contribute to some of the *de novo* glycine synthesis that we observed after *MTHFD2* or *SHMT2* knockdown; however, this is not sufficient to maintain elevated cellular glycine levels after TGF-β treatment. Our current and previous findings show that *SHMT1* and *MTHFD1* expression are decreased by TGF-β treatment (11), which may limit the ability of lung fibroblasts to utilize the reverse cytoplasmic 1C pathway in the absence of MTHFD2.

Our findings demonstrate that while the mitochondrial 1C pathway contributes to proline biosynthesis via NADP^+^ reduction, loss of MTHFD2 or ALDH1L2 is insufficient to decrease cellular proline levels downstream of TGF-β. While it would have been extremely elegant for the two most abundant amino acids in collagen to be linked in their mitochondrial production via an NADPH/NADP^+^ redox cycle, other sources of mitochondrial NADPH are sufficient to support proline production in the absence of one-carbon flux. NAD kinase 2 (NADK2) has recently been shown be required for proline synthesis (37, 38), and thus, phosphorylation of mitochondrial NAD(H) may be the primary mechanism by which NADPH is produced to support proline synthesis.

MTHFD2 is highly expressed during development, but exhibits low or absent expression in most adult tissues (39). *MTHFD2* expression is highly upregulated in a variety of cancers and is an independent prognostic indicator in breast and pancreatic cancer (39–41). Our findings demonstrate that MTHFD2 is highly expressed in fibrotic lungs and that increased *MTHFD2* expression correlates with loss of lung function in IPF patients. Due to the role of one-carbon metabolism in supporting cancer cell growth, inhibitors of this pathway are in development (42, 43). DS18561882 has been shown to inhibit tumor cell growth and T-cell mediated inflammation in mice (28–30). Our findings demonstrate that DS18561882 mimics the effect of *MTHFD2* knockdown on glycine accumulation and collagen production after TGF-β. Furthermore, DS18561882 was effective at inhibiting fibrotic responses after bleomycin instillation. While effects of DS18561882 on targets other than lung fibroblasts cannot be ruled out, MTHFD2 inhibition did not appear to have toxic effects on mice during the course of the experiment. Together, our findings suggest that MTHFD2 may be a viable target for fibrosis of the lung and other tissues.

## MATERIALS AND METHODS

### Fibroblast Culture

Normal human lung fibroblasts (NHLFs) (Lonza, CC-2512)) were maintained in Fibroblast Growth Medium 2 (PromoCell, C023020). Cells were plated at 1×10^5^ on 12 well plates for experiments. For experiments not involving metabolic labeling, 24 hours after plating, cells were serum starved in DMEM (Gibco, 11054020) supplemented with 0.1% bovine serum albumin and 2mM glutamine for 24 hours prior to treatment with 1ng/mL TGF-β (Peprotech, 100- 21C). DS18561882 (MedChemExpress, HY-130251) was added at the time of TGF-β treatment. For serine and glycine labeling experiments, MEM (Gibco, 11095080) was supplemented with MEM vitamin solution (Gibco, 1120052). For glycine labeling, 400μM ^13^C2 glycine (Cambridge Isotope Laboratories, CLM-1017) and 400μM unlabeled L-Serine (Sigma, 84959) were added. For serine labeling, 400μM 2,3,3-D3-L-Serine (Cambridge Isotope Laboratories, DLM-582) and 400μM unlabeled (Glycine Sigma, 50046).

### siRNA Knockdowns

For siRNA knockdowns, 1×10^6^ NHLFs were transfected with 250 pmol ON-TARGETplus siRNA (Dharmacon) using an Amaxa Nucleofector 2b set to program A-024. Cells were plated on 10cm dishes for 24 hours and then replated for experiments as above. Dharmacon product numbers: ATF4 (si1: J-005125-10, si2: J-005125-11), MTHFD2 (si1: J-009340-09, si2: J- 009349-11), MTHFD1L (si1: J-009949-09, si2: J-009949-12), ALDH1L2 (si1: J-026918-09, si2: J- 026918-12), MTHFD1 (si1: J-009577-05, si2: J-009577-06), SHMT2 (si1: J-004906-05, si2: J-004906-06).

### Western Blotting

Cells were lysed, and electrophoresis was performed as we previously described (44). Wells were lysed in 100μL Urea Sample Buffer (8M deionized urea, 1% SDS, 10% Glycerol, 60mM Tris pH 6.8, 0.1% pyronin-Y, 5% β-mercaptoethanol). Lysates were run through a 28 gauge needle and were electrophoresed on Criterion gels (Bio-Rad) and transferred to nitrocellulose using a Trans-Blot Turbo (Bio-Rad) set to the Mixed MW program. Primary antibodies used were: α-SMA (Sigma, A2547), ALDH1L2 (Proteintech, 21391-1-AP), ATF4 (Proteintech, 10835-1-AP), Collagen 1 (Abcam, ab138492), GAPDH (Cell Signaling, 2118), MTHFD1 (Proteintech, 10794-1-AP), MTHFD1L (Proteintech, 18112-1-AP), MTHFD2 (Proteintech, 12270-1-AP), Vinculin (Cell Signaling, 13901). Blots were imaged using a ChemiDoc Touch (Bio-Rad). Densitometry was performed using ImageJ.

### Quantitative PCR

RNA as isolated using GenElute Mammalian Total RNA Miniprep Kit (Sigma RTN350) and reverse transcribed using iScript Reverse Transcription Supermix (Bio-Rad, 1708841). Quantitative mRNA expression was determined by real-time RT-PCR using iTaq Universal SYBR Green Supermix (Bio-Rad, 172-5124). Primers used were: MTHFD2 (F: 5’- AAACACATCTGTCTGGTATGGT-3’, R: 5’-TGGTTAGGTCACAACTAGGAGTC-3’), ALDH1L2 (F: 5’-GCACTAATTGGCCAGAGCCT-3’, R: 5’-AGCCAGAGGGTCAGCTTTTC-3’), MTHFD1L (F: 5’-TTTGGTCGGAACGATGAGCA-3’, R: 5’-GTCCTGTGAGAGCCTTGTCC-3’), MTHFD1 (F: 5’- ACCCGGCCCTGTTTTTATGA-3’, R: 5’-TCCCAGTGGGCCTGAATAGA-3’), RPL13 (F: 5’- GTCGTACGCTGTGAAGGCAT-3’, R: 5’-GGAAAGCCAGGTACTTCAACTT-3’).

### Gas Chromatography-Mass Spectrometry

NHLFs grown for 48 hours in the presence or absence of TGF-β were washed in blood bank saline (Thermo, 23-293-184) and metabolites were extracted in 600μL ice cold 80% Methanol (Fisher, A456 (MeOH), W6 (H_2_O)). The solution was vortexed and centrifuged at 21,000 x g for 20 minutes. 400 μLof each extract was transferred to a new tube and dried under nitrogen. Dried metabolites were derivatized in 16μL Methoxamine reagent (Thermo, TS-45950) for 1hr at 37°C. 20 μL of 1% *N*-*tert*-butyldimethylsilyl-*N*- methyltrifluoroacetamide (Sigma, 394882) for 1hr at 60°C. Derivatized samples were analyzed with an 8890 gas chromatograph with an HP-5MS column (Agilent) coupled with a 5977B Mass Selective Detector mass spectrometer (Agilent). Helium was used as the carrier gas at a flow rate of 1.2ml/min. One microliter of each sample was injected in split mode (1:8) at 280°C. After injection, the GC oven was held at 100°C for 1 min and increased to 300°C at 3.5°C/min. The oven was then ramped to 320°C at 20°C/min and held for 5 minutes. The MS system was operated under electron impact ionization at 70eV and the MS source was operated at 230°C and quadrupole at 150°C. The detector was used in scanning mode, and the scanned ion range was 100-650 *m/z*. Peak ion chromatograms for metabolites of interest were extracted at their specific *m/z* with Mass Hunter Quantitative Analysis software (Agilent Technologies). Ions used for quantification of metabolite levels were as follows: Glycine *m/z* 246, Serine *m/z* 390, Proline *m/z* 258. Mass isotopomer distributions were determined by integrating appropriate ion fragments for each metabolite using IsoCor (45) to correct for natural abundance.

### Formate and Hydroxyproline Measurement

Formate content of media was measured using a Formate Assay Kit (Abcam, ab111748) on 50 μL of culture media following manufacturer’s instructions. For hydroxyproline measurements, 50 mg of cryogenically pulverized lung tissue was homogenized in 500 μL ultrapure H_2_O and hydroxy proline content was measured using a Hydroxyproline Assay Kit (Sigma-Aldrich, MAK357) following manufacturer’s instructions.

### Bleomycin-Induced Pulmonary Fibrosis

The protocol for the use of animals was approved by the University of Chicago Institutional Animal Care and Use Committee. Male C57Bl/6J mice (8- 12 weeks old; Jackson Laboratory) were intubated and bleomycin (0.7U/kg, TEVA Pharmaceuticals) was instilled intratracheally. Mice were sacrificed on the indicated days and perfused lung tissue was either snap frozen for hydroxyproline measurement or processed for histology. Mice receiving MTHFD2 inhibitor were IP injected with DS18561882 (MedChemExpress, HY-130251) beginning on day 8 after bleomycin instillation. Drug was administered at 100mg/kg in 150μL of 10% DMSO, 40% PEG300, 5% Tween-80, 45% saline.

### Micro Computed Tomography

MicroCT was performed on an X-Cube CT imager (Molecubes, Ghent, Belgium). Animals were anesthetized via inhalation of 2% isoflurane. MicroCT scan was performed using a respiratory-gated protocol to reduce motion artifacts. A window delay of 30% and window width of 50% was used to capture the end-expiration phase of the breathing cycle. Voltage was set to 50 kVp and current at 350 mAS. For quantitative 3D assessment of lung area, HU ranges [-900, −500] were considered as normally-aerated lung tissue, and HU ranges [-500, - 100] were defined as poorly-aerated. Volumetric microCT images were reconstructed in an 800 x 800 x 749 format with isotropic voxel dimensions of 50 x 50 x 50 μm3. Images were processed using VivoQuant 4 patch 1 (InviCRO).

### Immunohistochemistry

The collection and use of human lung specimens were approved by the University of Chicago Institutional Review Board. After deparaffinization and rehydration, tissue sections were treated with antigen retrieval buffer (DAKO, S1699) in a steamer for 20 min. Rabbit polyclonal anti-MTHFD2 antibody (Proteintech, 12270-1-AP, 1:400) was applied on tissue sections for a 1-h incubation at room temperature in a humidity chamber. Following a TBS wash, tissue sections were incubated with biotinylated anti-rabbit IgG (1:200, Vector Laboratories, BA- 1000) for 30 min at room temperature. The antigen-antibody binding was detected by Elite kit (PK-6100, Vector Laboratories) and DAB (DAKO, K3468) system. Tissue sections were briefly immersed in hematoxylin for counterstaining and were covered with cover glasses. Whole slide images were analyzed using ImageJ.

### Analysis of gene expression and patient data

Processed gene expression datasets from GSE32537 were downloaded from Gene Expression Omnibus. Available clinical data was correlated to individual gene expression profiles using Pearson’s correlation analysis using GraphPad Prism v10.0.2.

### Statistical analysis

Data were analyzed in Prism 10 (GraphPad Software, Inc). All data are shown as mean ± standard error of the mean (SEM). Significance was determined by unpaired two-tailed Student’s t test (for comparisons between two samples), or by one or two-way ANOVA using Tukey’s multiple comparison test. * P < 0.05, ** P < 0.01, *** P < 0.001.

## SUPPLEMENTARY FIGURE LEGENDS

**Figure S1.**
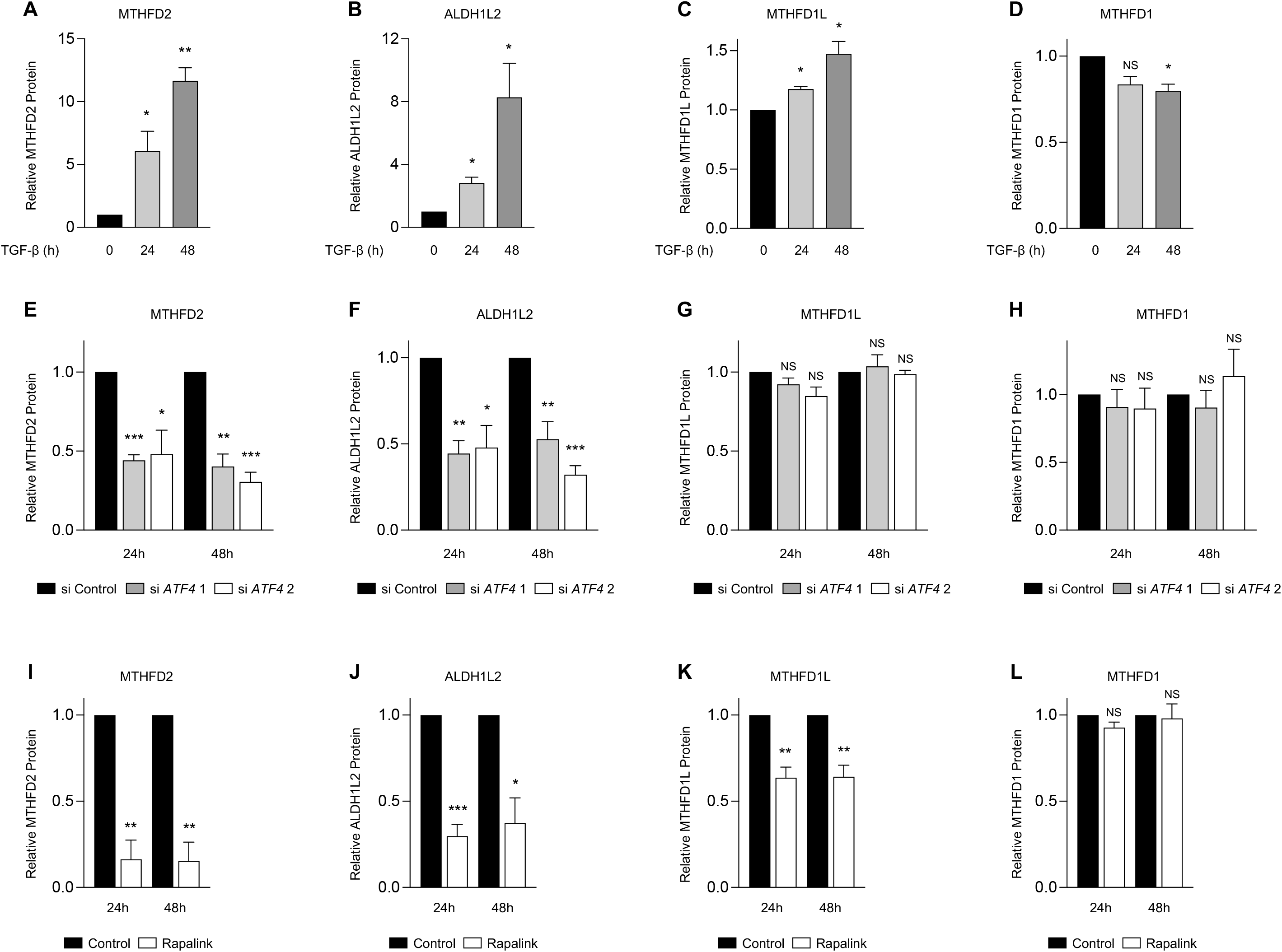
Mitochondrial one-carbon enzymes are induced by TGF-β. **(A-D)** Quantification of relative (A) MTHFD2, (B) ALDH1L2, (C) MTHFD1L, and (D) MTHFD1 protein levels after treatment of normal human lung fibroblasts (NHLFs) treated with TGF-β for the indicated intervals. **(E-H)** Quantification of relative (E) MTHFD2, (F) ALDH1L2, (G) MTHFD1L, and (H) MTHFD1 protein levels after treatment of NHLFs transfected with siRNA targeting ATF4 or nontargeting siRNA. Cells were treated with TGF-β for the indicated intervals. **(I-L)** Quantification of relative (I) MTHFD2, (J) ALDH1L2, (K) MTHFD1L, and (L) MTHFD1 protein levels after treatment of NHLFs with TGF-β for the indicated intervals in the presence or absence of Rapalink-1. Bar graphs represent mean ± SEM, n=3 independent experimental replicates. \**P*<0.05, \*\**P*<0.01, \*\*\**P*<0.001.

**Figure S2.**
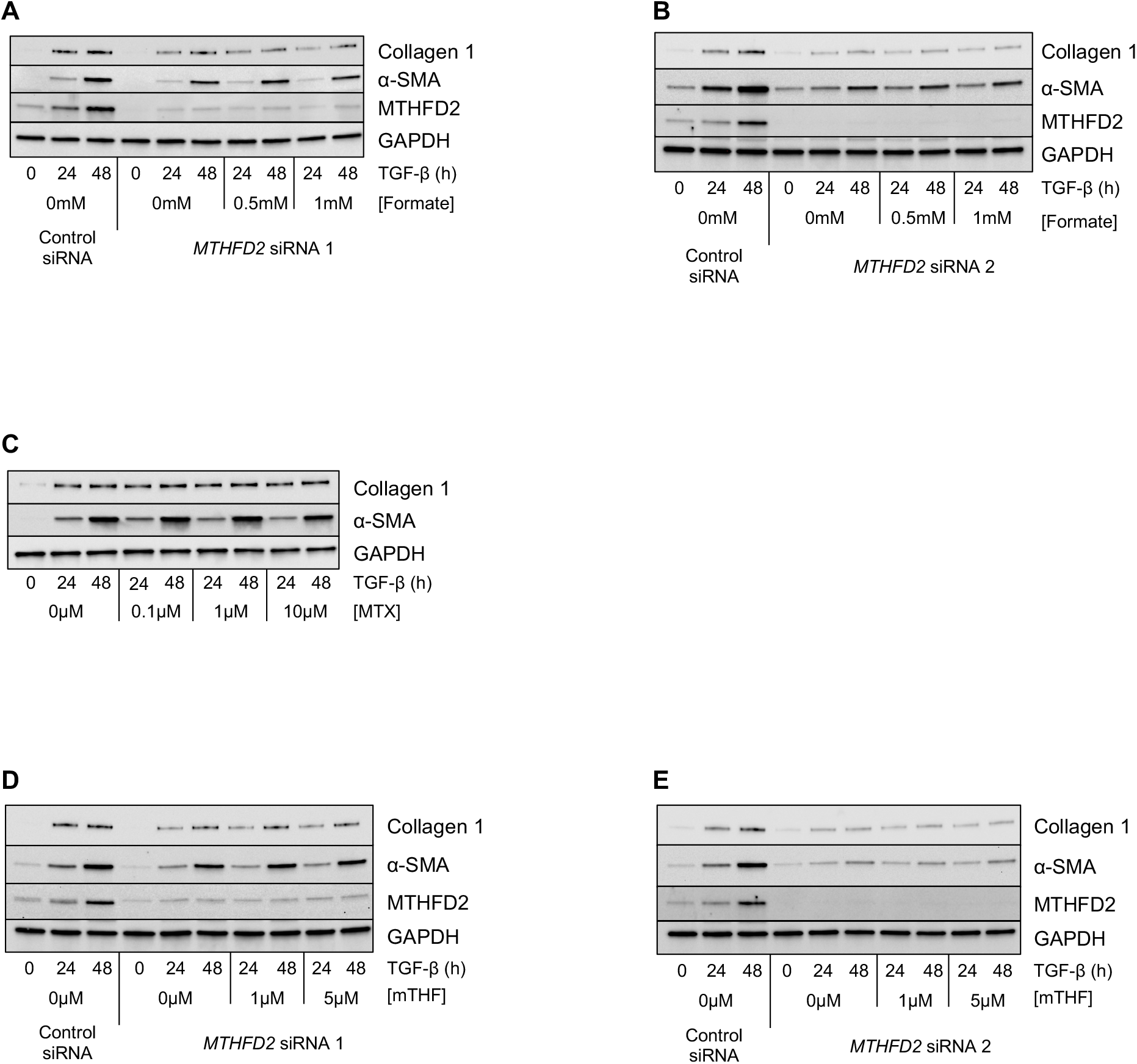
Mitochondrial one-carbon metabolism is required for TGF-β-induced collagen protein production. **(A-B)** Western blot analysis of collagen and α-smooth muscle actin (α-SMA) protein expression in normal human lung fibroblasts (NHLFs) transfected with siRNAs targeting *MTHFD2* or nontargeting siRNA. Cells were treated with TGF-β for the indicated intervals in the presence of the indicated doses of formate. **(C)** Western blot analysis of collagen and α-SMA protein expression in NHLFs treated TGF-β for the indicated intervals in the presence of the indicated doses of methotrexate (MTX). **(D-E)** Western blot analysis of collagen and α-SMA protein expression in NHLFs transfected with siRNAs targeting *MTHFD2* or nontargeting siRNA. Cells were treated with TGF-β for the indicated intervals in the presence of the indicated doses of 5-methyl-tetrahydrofolate (mTHF).

**Figure S3.**
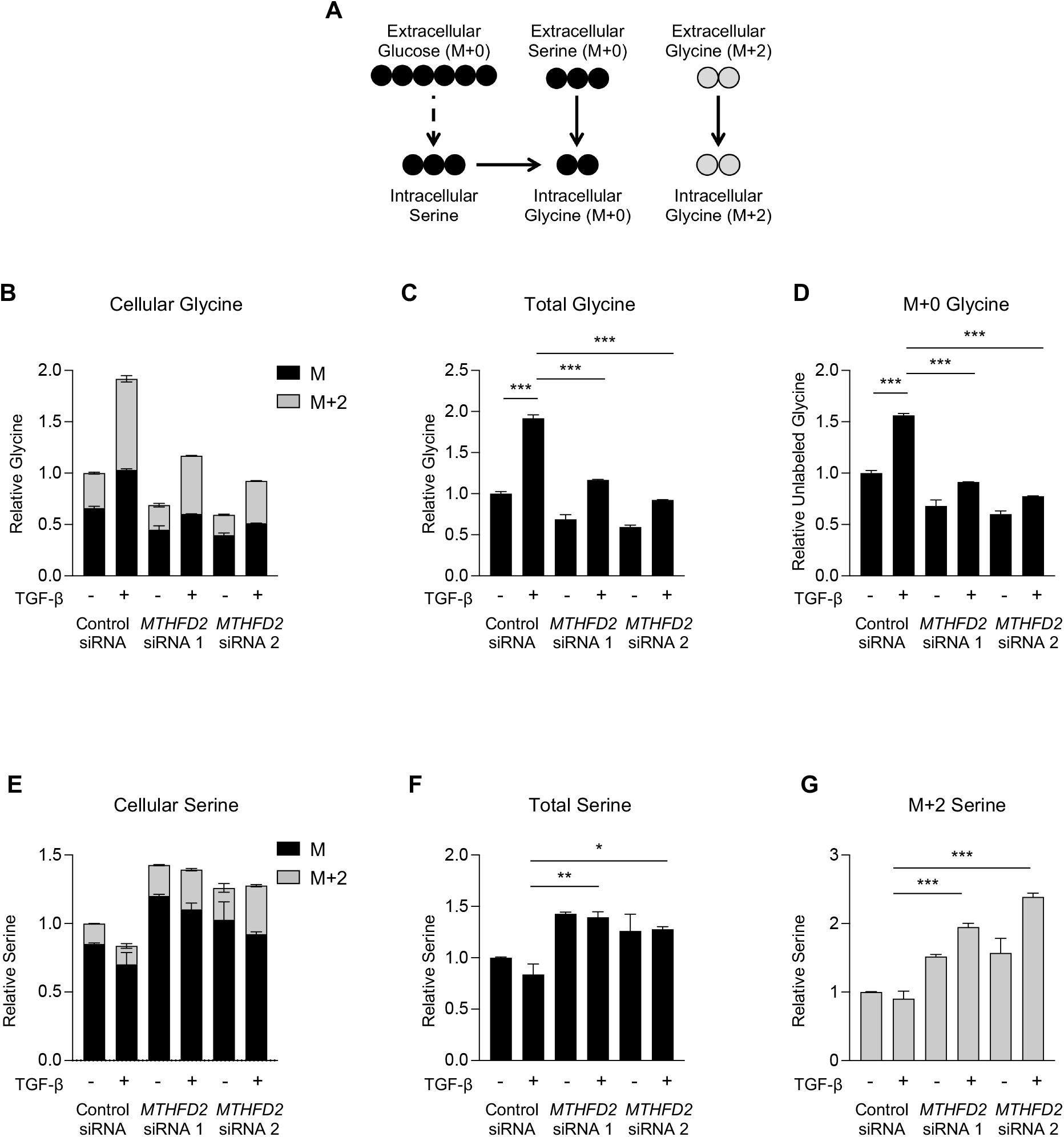
MTHFD2 is required for increased cellular glycine levels downstream of TGF-β. **(A)** Schematic representation of metabolite labeling downstream of ^13^C2-Glycine. Normal human lung fibroblasts (NHLFs) were transfected with siRNA targeting *MTHFD2* or nontargeting siRNA. Cells were labeled with ^13^C2-Glycine and treated with TGF-β for 48 hours or left untreated. **(B)** Analysis of cellular glycine after labeling with ^13^C2-Glycine in NHLFs treated with TGF-β or left untreated. **(C)** Relative total glycine levels from (B). **(D)** Relative levels of M+0 (*de novo* synthesized) glycine from (B). **(E)** Analysis of cellular serine after labeling with ^13^C2-Glycine in NHLFs treated with TGF-β or left untreated. **(F)** Relative total serine levels from (E). **(G)** Relative levels of M+2 serine from (E). Bar graphs represent mean ± SEM, n=3. \**P*<0.05, \*\**P*<0.01, \*\*\**P*<0.001.

**Figure S4.**
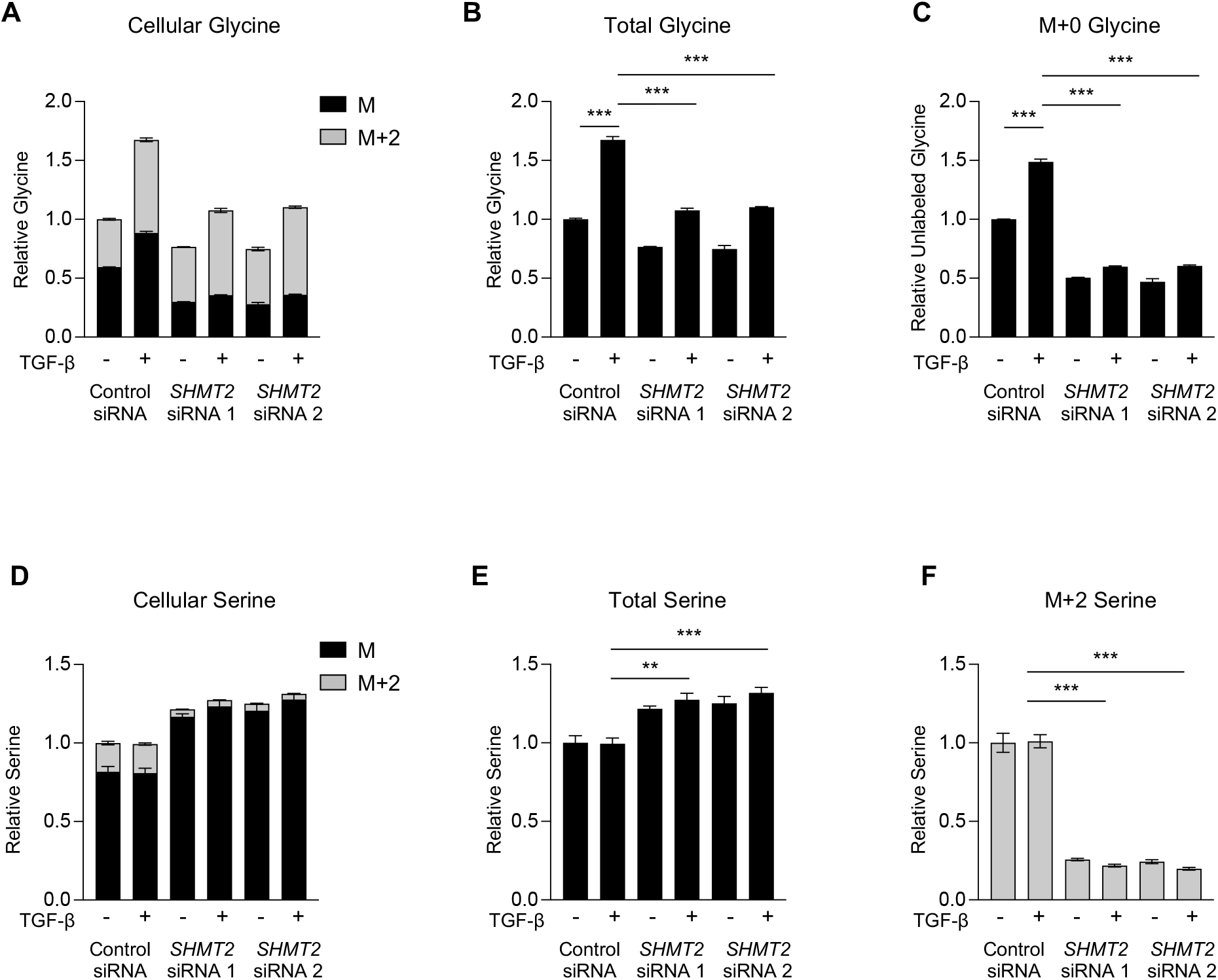
SHMT2 is required for increased cellular glycine levels downstream of TGF-β. Normal human lung fibroblasts (NHLFs) were transfected with siRNA targeting *SHMT2* or nontargeting siRNA. Cells were labeled with ^13^C2-Glycine and treated with TGF-β for 48 hours or left untreated. **(A)** Analysis of cellular glycine after labeling with ^13^C2-Glycine in NHLFs treated with TGF-β or left untreated. **(B)** Relative total glycine levels from (A). **(C)** Relative levels of M+0 (*de novo* synthesized) glycine from (A). **(D)** Analysis of cellular serine after labeling with ^13^C2- Glycine in NHLFs treated with TGF-β or left untreated. **(E)** Relative total serine levels from (D). **(F)** Relative levels of M+2 serine from (D). Bar graphs represent mean ± SEM, n=3. \**P*<0.05, \*\**P*<0.01, \*\*\**P*<0.001.

**Figure S5.**
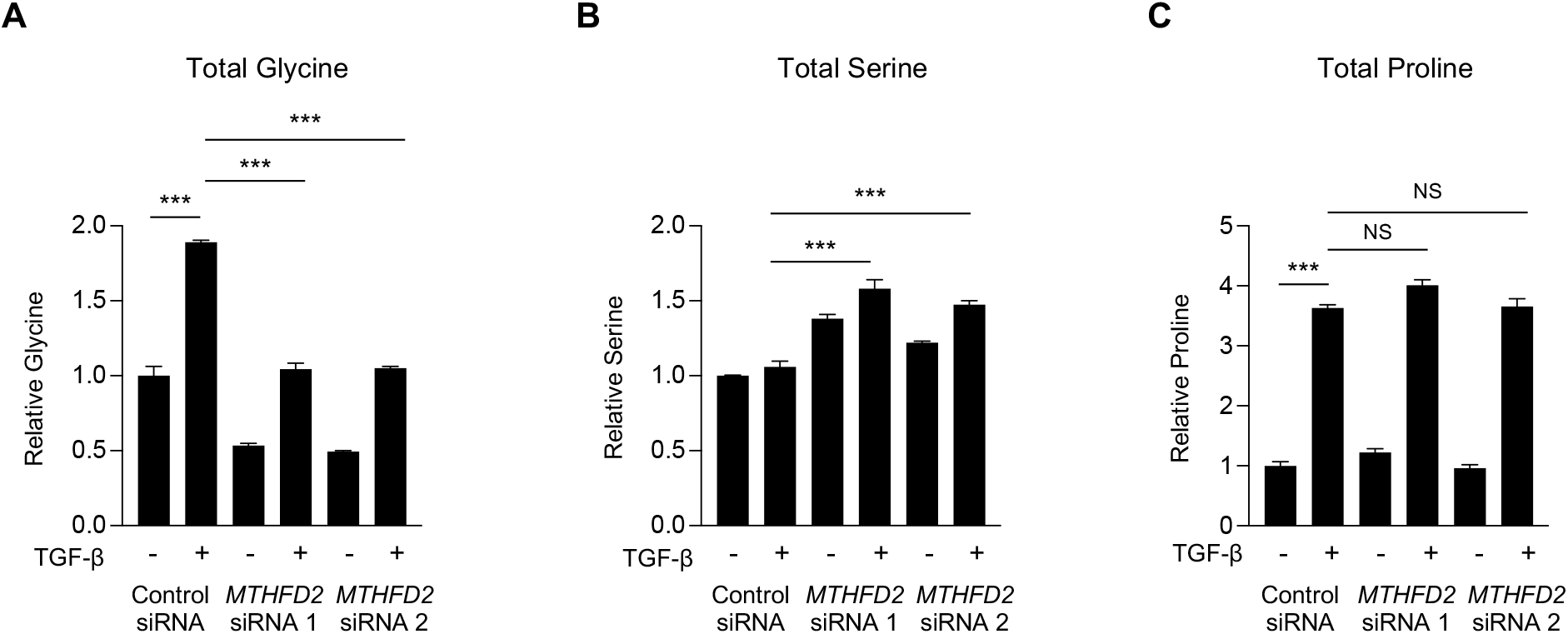
Analysis of cellular metabolite levels in *MTHFD2* knockdown human lung fibroblasts. Normal human lung fibroblasts (NHLFs) were transfected with siRNA targeting *MTHFD2* or nontargeting siRNA. Cells were labeled with 2,3,3-D3-Serine and treated with TGF- β for 48 hours or left untreated. **(A)** Relative total glycine levels from Figure 3B. **(B)** Relative total serine levels from Figure 3D. **(C)** Relative total proline levels from Figure 3F. Bar graphs represent mean ± SEM, n=3. \**P*<0.05, \*\**P*<0.01, \*\*\**P*<0.001.

**Figure S6.**
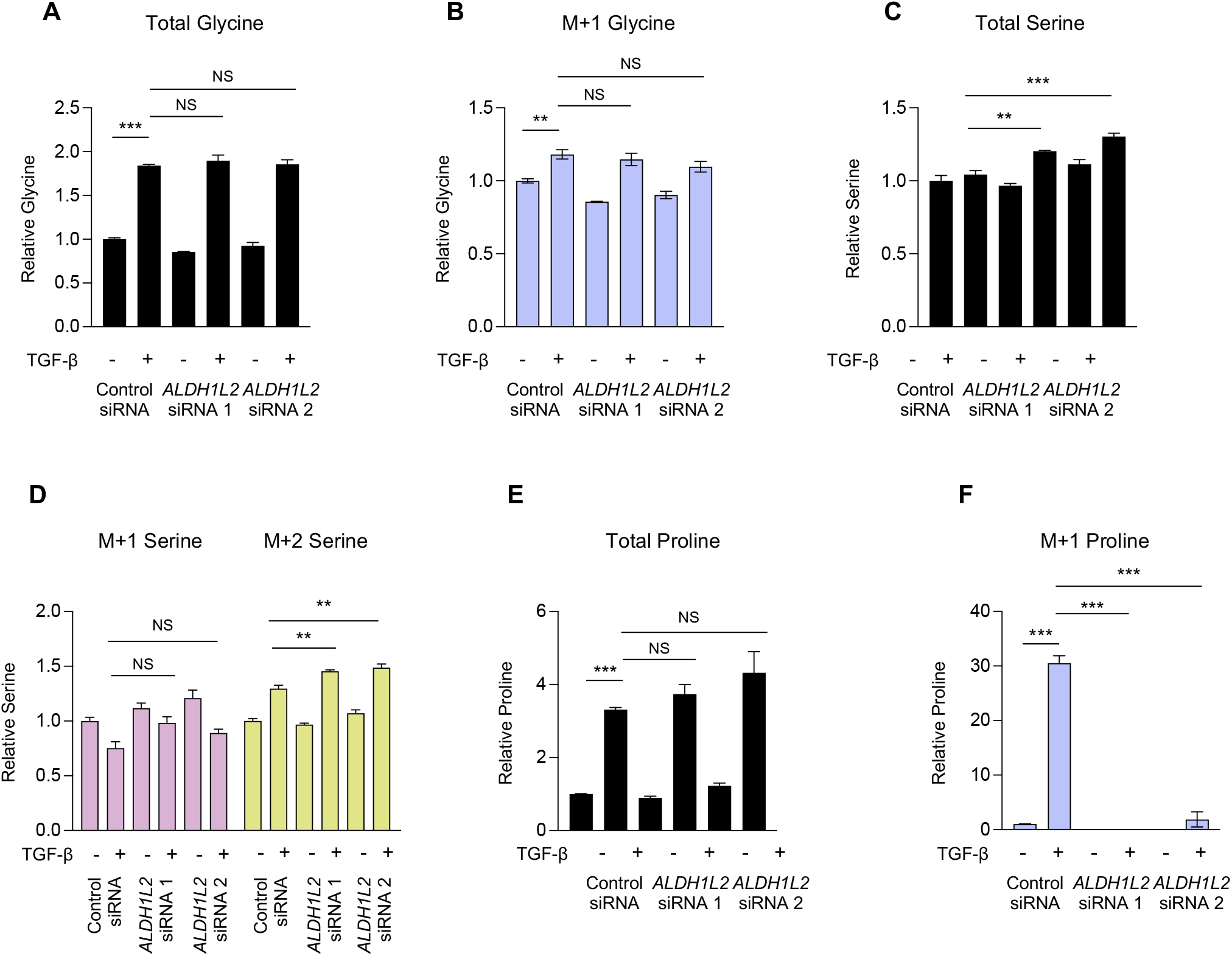
Analysis of cellular metabolite levels in *ALDH1L2* knockdown human lung fibroblasts. Normal human lung fibroblasts (NHLFs) were transfected with siRNA targeting *ALDH1L2* or nontargeting siRNA. Cells were labeled with 2,3,3-D3-Serine and treated with TGF- β for 48 hours or left untreated. **(A)** Relative total glycine levels from Figure 4A. **(B)** Relative levels of M+1 glycine from Figure 4A. **(C)** Relative total serine levels from Figure 4B. **(D)** Relative levels of M+1 and M+2 serine from Figure 4B. **(E)** Relative total proline levels from Figure 4C. **(F)** Relative levels of M+1 proline from Figure 4C. Bar graphs represent mean ± SEM, n=3. \**P*<0.05, \*\**P*<0.01, \*\*\**P*<0.001.

**Figure S7.**
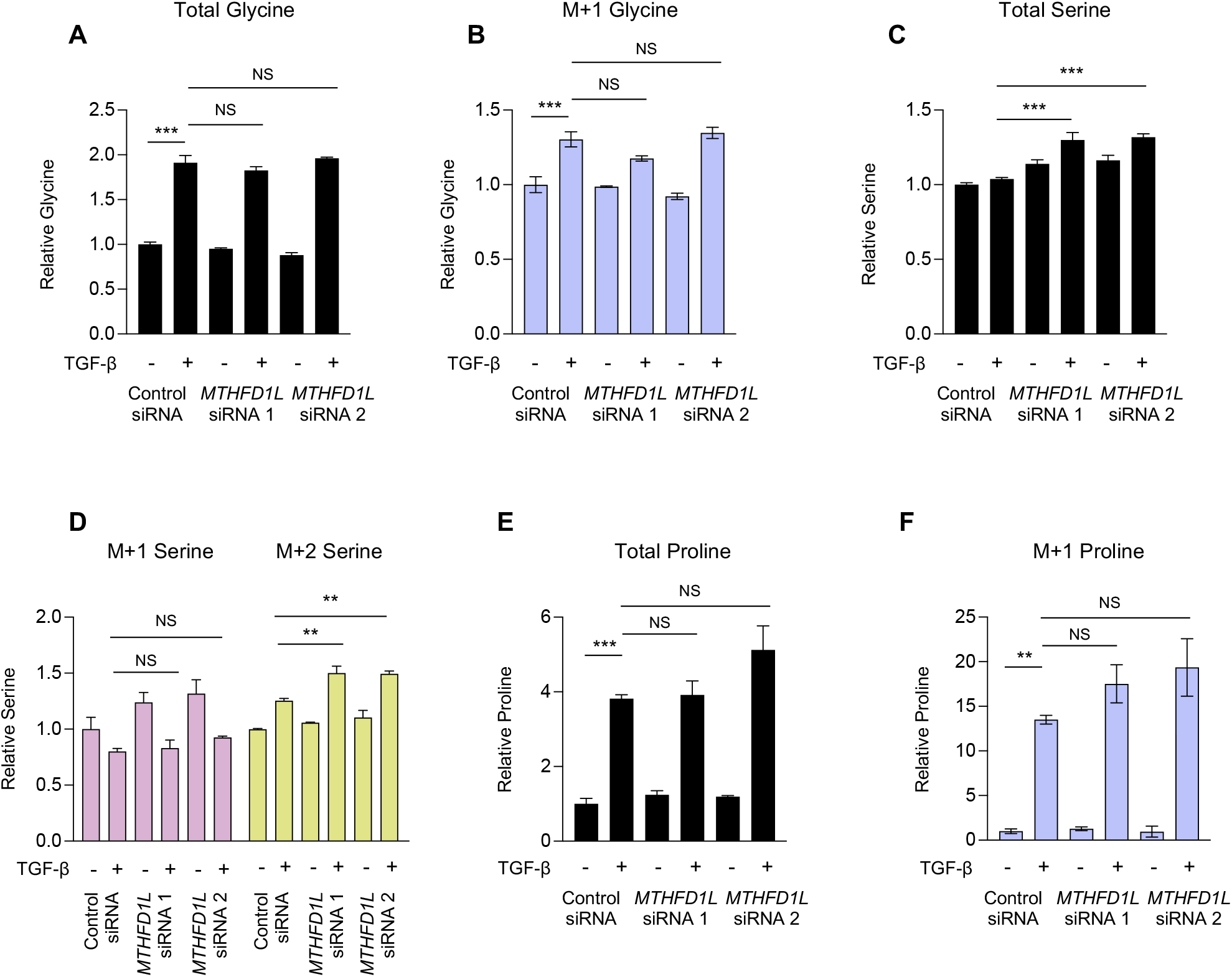
Analysis of cellular metabolite levels in *MTHFD1L* knockdown human lung fibroblasts. Normal human lung fibroblasts (NHLFs) were transfected with siRNA targeting *MTHFD1L* or nontargeting siRNA. Cells were labeled with 2,3,3-D3-Serine and treated with TGF- β for 48 hours or left untreated. **(A)** Relative total glycine levels from Figure 4E. **(B)** Relative levels of M+1 glycine from Figure 4E. **(C)** Relative total serine levels from Figure 4F. **(D)** Relative levels of M+1 and M+2 serine from Figure 4F. **(E)** Relative total proline levels from Figure 4G. **(F)** Relative levels of M+1 proline from Figure 4G. Bar graphs represent mean ± SEM, n=3. \**P*<0.05, \*\**P*<0.01, \*\*\**P*<0.001.

**Figure S8.**
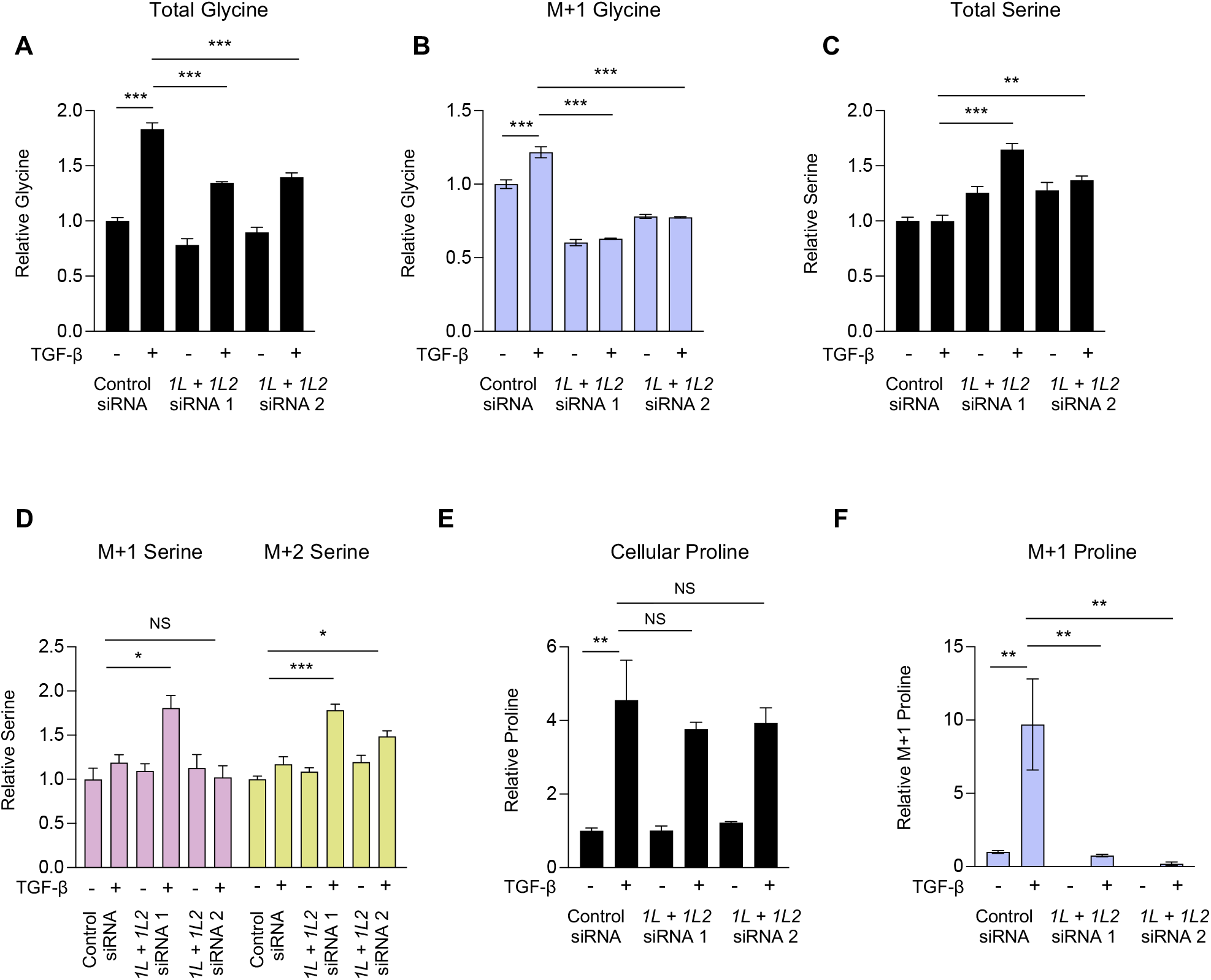
Analysis of cellular metabolite levels in human lung fibroblasts with knockdown of both *ALDH1L2* and *MTHFD1L*. Normal human lung fibroblasts (NHLFs) were transfected with siRNA targeting both *ALDH1L2* and *MTHFD1L* or nontargeting siRNA. Cells were labeled with 2,3,3-D3-Serine and treated with TGF-β for 48 hours or left untreated. **(A)** Relative total glycine levels from Figure 4I. **(B)** Relative levels of M+1 glycine from Figure 4I. **(C)** Relative total serine levels from Figure 4J. **(D)** Relative levels of M+1 and M+2 serine from Figure 4J. **(E)** Relative total proline levels from Figure 4K. **(F)** Relative levels of M+1 proline from Figure 4K. Bar graphs represent mean ± SEM, n=3. \**P*<0.05, \*\**P*<0.01, \*\*\**P*<0.001.

**Figure S9.**
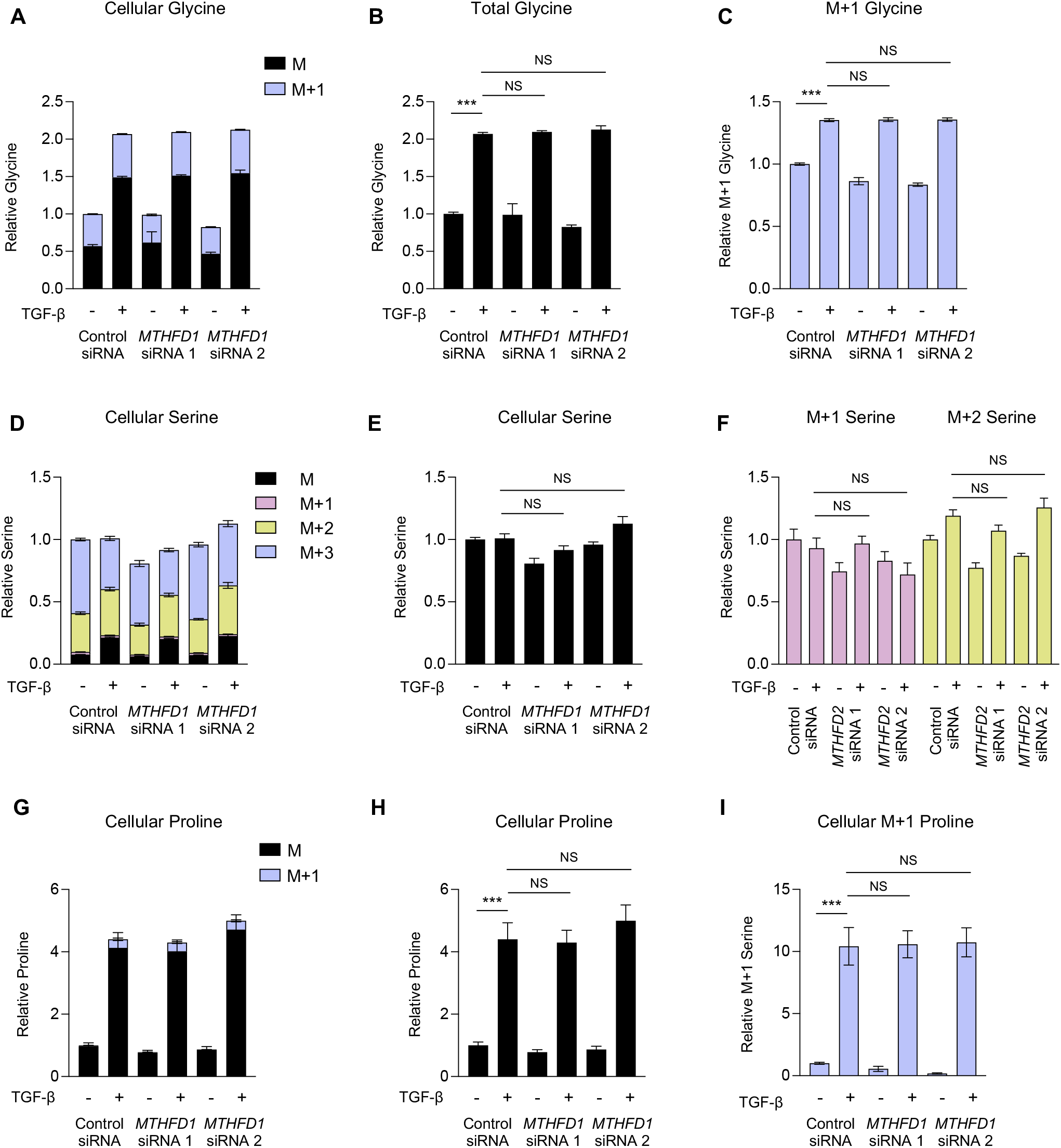
*MTHFD1* is not required for increased cellular glycine levels downstream of TGF-β. Normal human lung fibroblasts (NHLFs) were transfected with siRNA targeting *MTHFD1* or nontargeting siRNA. Cells were labeled with 2,3,3-D3-Serine and treated with TGF-β for 48 hours or left untreated. **(A)** Analysis of cellular glycine after labeling with 2,3,3-D3-Serine in NHLFs treated with TGF-β or left untreated. **(B)** Relative total glycine levels from (A). **(C)** Relative levels of M+1 glycine from (A). **(D)** Analysis of cellular serine after labeling with 2,3,3-D3-Serine in NHLFs treated with TGF-β or left untreated. **(E)** Relative total serine levels from (D). **(F)** Relative levels of M+1 and M+2 serine from (D). **(G)** Analysis of cellular proline after labeling with 2,3,3- D3-Serine in NHLFs treated with TGF-β or left untreated. **(H)** Relative total serine levels from (G). **(I)** Relative levels of M+1 and M+2 serine from (G). Bar graphs represent mean ± SEM, n=3. \**P*<0.05, \*\**P*<0.01, \*\*\**P*<0.001.

**Figure S10.**
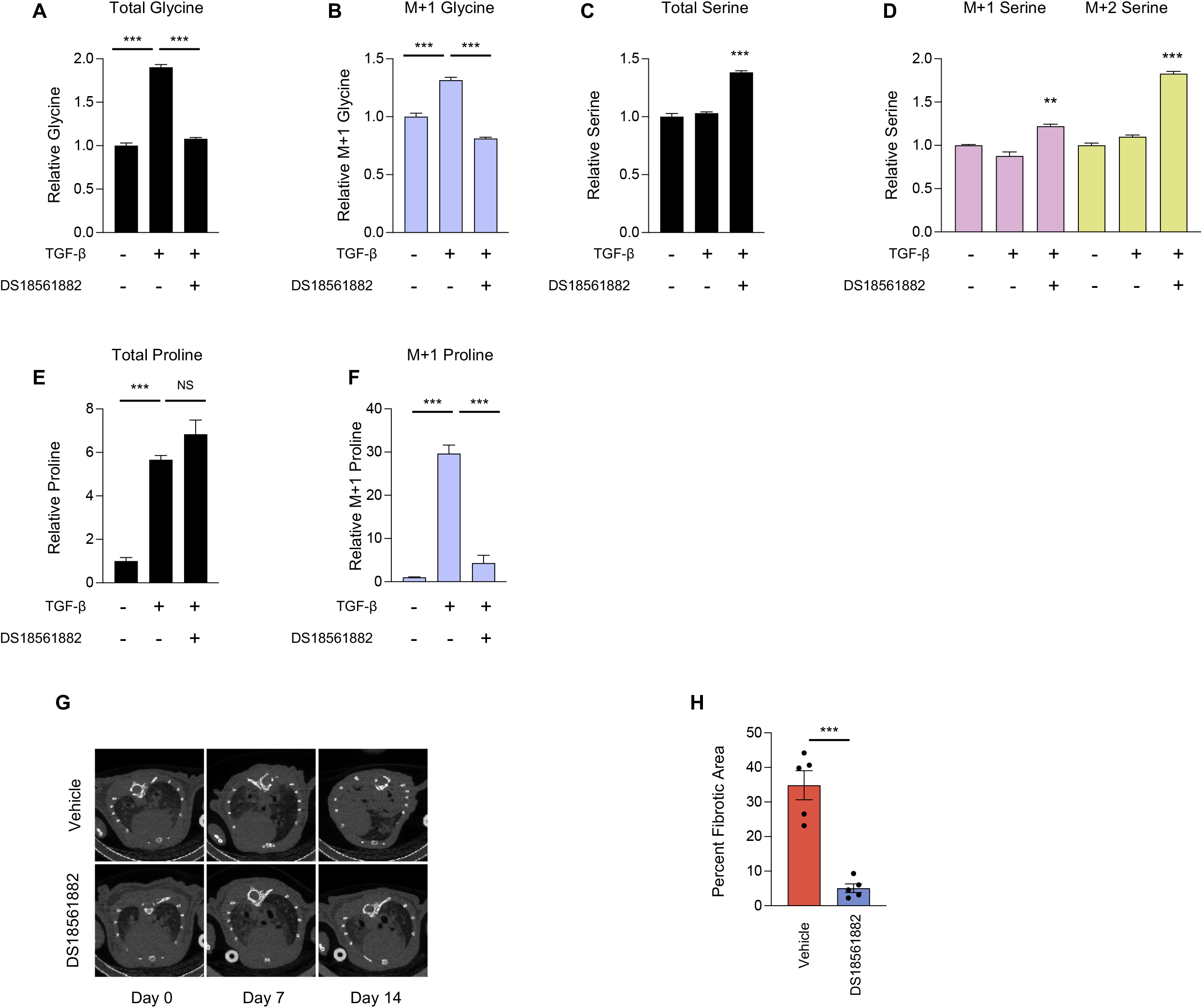
Pharmacologic inhibition of MTHFD2 inhibits fibrotic responses. **(A)** Relative total glycine levels from Figure 5E. **(B)** Relative levels of M+1 glycine from Figure 5E. **(C)** Relative total serine levels from Figure 5F. **(D)** Relative levels of M+1 and M+2 serine from Figure 5F. **(E)** Relative total proline levels from Figure 5G. **(F)** Relative levels of M+1 proline from Figure 5G. Bar graphs represent mean ± SEM, n=3. **(G)** Representative transverse microCT images of mice prior to bleomycin instillation (Day 0), 7 days after bleomycin instillation (Day 7), and 14 days after bleomycin instillation (Day 14). Mice began receiving DS18561882 (125 mg/kg) or vehicle 8 days after bleomycin instillation. **(H)** Percent fibrotic area of lung from mice 21 days after bleomycin instillation. Mice received either vehicle or DS18561882 beginning on day 8. Bar graphs represent mean ± SEM, n=5 independent mice per condition. \**P*<0.05, \*\**P*<0.01, \*\*\**P*<0.001.

